# Investigating pedigree- and SNP-associated components of heritability in a wild population of Soay sheep

**DOI:** 10.1101/2023.06.02.543397

**Authors:** Caelinn James, Josephine M. Pemberton, Pau Navarro, Sara Knott

## Abstract

Estimates of narrow sense heritability derived from genomic data that contain related individuals may be biased due to the within-family effects such as dominance, epistasis and common environmental factors. However, for many wild populations, removal of related individuals from the data would result in small sample sizes. In 2013, Zaitlen et al. proposed a method to estimate heritability in populations that include close relatives by simultaneously fitting an identity-by-state genomic relatedness matrix (GRM) and an identity-by-descent GRM. The IBD GRM is identical to the IBS GRM, except relatedness estimates below a specified threshold are set to 0. We applied this method to a sample of 8557 wild Soay sheep from St. Kilda, with genotypic information for 419,281 single nucleotide polymorphisms to investigate polygenic and monogenic traits. We also implemented a variant of the model in which the IBD GRM was replaced by a GRM constructed from SNPs with low minor allele frequency to examine whether any additive genetic variance is captured by rare alleles. Each model was compared to an animal model with a single GRM based on all genotyped markers (the IBS GRM) using a log likelihood ratio test. Whilst the inclusion of the IBD GRM did not significantly improve the fit of the model for the monogenic traits, it improved the fit for some of the polygenic traits, suggesting that dominance, epistasis and/or common environment not already captured by the non-genetic random effects fitted in our models may influence these traits.

## Introduction

The “genetic architecture” of a trait is a broad term used to describe the characteristics of genetic variants that contribute to the trait’s phenotypic variation. Such characteristics include the number of loci contributing towards variation, the amount of variation attributable to each causal locus, the location of these loci in the genome, and how rare or common causal alleles are in the population. It is important to understand the genetic architecture underpinning traits of interest; in disease research it can be used to inform clinical diagnosis and prognosis as well as identify potential treatments; in livestock and crop breeding, it allows for more informed selective breeding strategies to improve trait yield; in evolutionary genetics we can use knowledge of the genetic architecture of a trait to understand how evolutionary processes may act on a trait and what micro-evolutionary dynamics are occurring in the population.

Historically, heritability estimation has been a key metric when investigating the genetic architecture of a trait. Narrow sense heritability (h^2^) is the proportion of phenotypic variation which is explained by additive genetic variation in the population, and traditionally was estimated using correlations between family members in twin studies or through parent-offspring regression. More recently the animal model (Meyer 1989) has become widely used, and has the advantage of using relationships between all individuals and across generations to increase power and better estimate the additive genetic component. The additive genetic variance can be estimated by an animal model using relatedness inferred from either the pedigree (h^2^_ped_ – obtained using the numerator relationship matrix A (Henderson 1975)) or genetic markers such as SNPs (h^2^_GRM_ – obtained using a genomic relationship matrix (GRM) (Yang et al. 2010)). The pedigree captures variance due to identity-by-descent and provides estimates of expected relatedness rather than realized relatedness. The pedigree can also be prone to error, especially when it is derived from observational data, but also in cases when it is derived from genotype data but paternities cannot be accurately or uniquely assigned. A conventional GRM captures identity-by-state, and the resulting estimates are dependent on which SNPs are used to calculate the GRM; if neither the causal SNP nor any SNPs in linkage disequilibrium (LD) with the causal SNP have been genotyped, the h^2^_GRM_ estimate may be underestimated. This can be avoided by using high density genotyping arrays or full genome sequencing, however in some wild populations it has been shown that increasing the number of SNPs above a certain threshold does not affect GRM-based heritability estimates (Bérénos et al. 2014; Perrier et al. 2018; James et al. 2022).

Inclusion of related individuals when estimating h^2^_GRM_ can result in overestimation by inadvertently capturing effects of common environment, epistasis and dominance. However, removal of related individuals is not optimal for smaller study populations such as wild populations, as this reduces sample sizes. In 2013, Zaitlen et al. proposed a partitioning method that simultaneously estimates both h^2^ and h^2^_GRM_ (referred to as h^2^_g_ by the authors) of a trait in a population that contains related individuals. The method involves running an animal model simultaneously fitting both a GRM and an additional GRM which is thresholded in such a way that relatedness estimates below a specified threshold are set to 0. Both GRMs jointly model variance due to identity-by-descent (IBD), whilst the non-thresholded GRM models variance due to identity-by-state (IBS). This method has so far been applied to a range of traits in humans (Xia et al. 2016; Hill et al. 2018) as well as in wild passerine birds (Silva et al. 2017) and farmed salmon (Kokkinias 2022). The extent to which thresholded GRMs explain trait variance differs across these studies, presumably due to differences in relatedness structure, trait architecture, trait heritability and other variables, so it is of interest to explore the outcome of the approach in a wide range of populations and traits.

Here, we use the method of Zaitlen et al. (2013) to simultaneously estimate h^2^ and h^2^_GRM_ for a selection of polygenic and monogenic traits in a wild population of related Soay sheep. The effect of additive genetic variation segregating at the population level is fitted using the non-thresholded IBS GRM, whilst the family-associated variation (such as epistasis, dominance, common environmental factors not captured by our non-genetic random effects fitted in the model, and any additional additive genetic variation segregating within families) is captured using the thresholded IBD GRM. In addition we explored a modified version of the approach, in which the thresholded IBD GRM was replaced with a GRM calculated from SNPs that had a minor allele frequency (MAF) below a certain threshold (but not thresholding relatedness), allowing us to examine how much of the genetic variation is captured by rare alleles and whether a rare-allele GRM can be used as a substitute for a relatedness thresholded GRM. We analysed a mixture of monogenic and polygenic traits in order to better understand how extremes of the genetic architecture of a trait affect the outcome of our heritability partition.

## Methods

### Phenotypic data

The Soay sheep (*Ovis aries*) of the St. Kilda archipelago is a primitive breed of sheep that has been the focus of a longitudinal, individual-based study since 1985 (Clutton-Brock and Pemberton 2003). Individuals are ear-tagged at first capture (usually two to ten days after birth) to allow re-identification, regularly recaptured in order to measure, amongst others, various morphometric and life history traits across an individual’s lifespan, and, after death, skeletal remains are collected and measured.

We focused on five polygenic morphometric traits across three different age classes (neonate, lamb and adult). Birth weight was measured in neonates. Three traits (weight, foreleg length and hindleg length) were measured on live individuals during an August catch up for lambs and adults. As adults are often recaptured across different years, the live traits have repeated measurements for many individuals. The remaining two traits (metacarpal length and jaw length) are *post mortem* measures taken from skeletal remains. Both birth and August weight are measured to the nearest 0.1kg, whilst the remaining traits are all measured to the nearest mm. A detailed description of trait measurements can be found in Beraldi et al. (2007). Several studies have demonstrated that these traits are under polygenic control (Bérénos et al. 2015; Ashraf et al. 2021; Hunter et al. 2022; James et al. 2022).

Neonates were defined as individuals who were caught and weighed between two and ten days after birth. For live traits, lambs were defined as individuals whose morphometric data was recorded in the August of their birth year, and for *post mortem* traits in lambs were individuals who died before 14 months of age. Similarly, adults were defined as individuals with phenotypic data recorded at least two years after birth for the live traits, or if they died after 26 months of age for the *post mortem* traits. We did not include yearling data in our analyses due to the small sample sizes available following first winter mortality.

In addition to the polygenic traits, we analysed four monogenic traits (male horn type, female horn type, coat colour and wild/self coat pattern) to examine whether the different genetic architectures underpinning polygenic and monogenic traits yield different results when partitioning the heritability. These traits are well characterized in this population and the causal gene for each trait has been identified (see Johnston et al. (2013), Gratten et al. (2007), Gratten et al. (2010) and Supplementary Table 1 for additional information on the underlying genetics of horn type, coat colour and coat pattern respectively). Male and female horn types were analysed separately as while both are controlled by the same locus, in males the “normal horned” allele is dominant, so only two phenotype classes (normal horned and scurred) are observed, whilst in females there is codominance and three phenotypes are observed (normal horned, scurred and polled). As the monogenic traits investigated here do not change over ontogeny, we did not analyse these traits by age class.

Table 1 lists the number of individuals and records per trait.

### Genetic data

Data were available for 8557 sheep genotyped on the Ovine SNP50 Illumina BeadChip, which gave genotypes for 38,130 autosomal variants that are polymorphic in the population. A subset of 438 SNPs are used to recover the pedigree using Sequoia (Huisman 2017). 188 individuals have also been genotyped on the Ovine Infinium HD SNP BeadChip, which has a much higher density of variants than the SNP50 BeadChip. This has allowed for the remaining genotypes to be imputed to this higher density using AlphaImpute, which combines shared haplotype and pedigree information for phasing and imputation (Hickey et al. 2012) (see Stoffel et al. (2021) for information on our imputation procedure). We used imputed genotype “hard” calls rather than genotype probabilities in downstream analyses. After filtering SNPs that passed quality control standards, 419,281 autosomal SNPs remained for 8557 individuals (4035 females, 4452 males).

### Variance component analysis

We used animal models to partition the phenotypic variance for each trait into genetic and non-genetic variance components. For each trait, we ran multiple models:

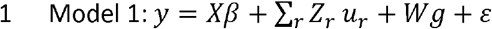

Where **y** is the vector of phenotypic values; **X** is a design matrix linking individual records with the vector of fixed effects **β, Z**_**r**_ is an incidence matrix that relates a non-genetic random effect to the individual records; **u**_**r**_ is the associated vector of non-genetic random effects; **g** is the vector of additive genetic random effects of the whole study population with **W** the incidence matrix; and **ε** is the vector of residuals. It is assumed that **g** ∼ *MVN*(0, M_g_σ_g_^2^), where σ_g_^2^ is the additive genetic variance and M_g_ is the GRM for the whole study population. This model was run once per trait for the polygenic traits. For the monogenic traits, this model was run three times: once with all SNPs being used to construct the GRM, once with all SNPs on the same chromosome as the causal gene being used to construct the GRM, and once with all SNPs within 1Mb upstream or downstream of the causal gene being used to construct the GRM. For monogenic traits, these models were designed to eliminate noise from genetic variation from non-causal loci, given that the only variants affecting these traits should be those within the previously identified causal regions.

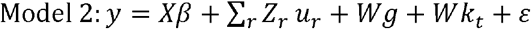

The terms in this model are the same as in Model 1, with the inclusion of **k**_**t**_, the vector of extra genetic random effects associated with relatives with a genomic relatedness higher than threshold t. It is assumed that **k**_**t**_ ∼ *MVN*(0, M_kt_σ_kt_^2^), where σ_kt_^2^ is the kinship genetic variance and M_kt_ is the kinship GRM with relationships equal to or less than t being set to 0. This model was run twice per trait, once at t=0.05 and once at t=0.1. Both thresholds capture parent-offspring, full sibling and half-sibling relationships but differ as to the proportion of more distantly related pairs of individuals that are retained.

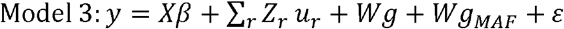

The terms in this model are the same as in Model 1, with the inclusion of **g**_**MAF**_, the vector of additive genetic random effects of SNPs with a MAF under a specified threshold in the whole study population. It is assumed that **g**_**MAF**_ ∼ *MVN*(0, M_MAF_σ_MAF_^2^), where σ_MAF_^2^ is the additive genetic variance of SNPs with a MAF below a set threshold, and M_MAF_ is the GRM calculated from the SNPs with a MAF below the threshold for the whole study population and range of relationships. For polygenic traits, this model was run five times, with MAF thresholds varying between 0.1 and 0.001 (see Table 2 for the full list of MAF thresholds and the number of SNPs that remained for each threshold).

For the monogenic traits, in addition to running the same models as for the polygenic traits, we also ran the model for each threshold where both GRMs were only constructed from SNPs on the chromosome containing the causal gene, and again for each threshold where both GRMs were only constructed from SNPs within the region spanning 1Mb either side of the causal gene (See Table 3 for the number of SNPs that remained for each threshold for each trait for both the single chromosome and region based GRMs). We did not run the model for any thresholds that resulted in a GRM being computed for less than 10 SNPs. These models were designed to eliminate noise from genetic variation from non-causal loci, given that the only variants affecting these traits should be those within the previously identified causal regions.

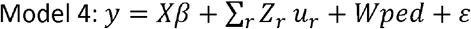

The terms in this model are the same as in Model 1, with the exception of **ped** being the vector of pedigree based effects of the whole study population. It is assumed that **ped** ∼ *MVN*(0, A_g_σ_g_^2^), where σg^2^ is the additive genetic variance and A_g_ is the relationship matrix for the whole study population. Again, this model was run once per trait.

Fixed and non-genetic random effects were only fitted for the polygenic traits, with the effects fitted differing between traits and age classes (see Table 1). For each trait, the same individuals were analysed across all models.

The GRMs were computed using GCTA (Yang et al. 2011), which computes the genetic relationship between individuals i and j as

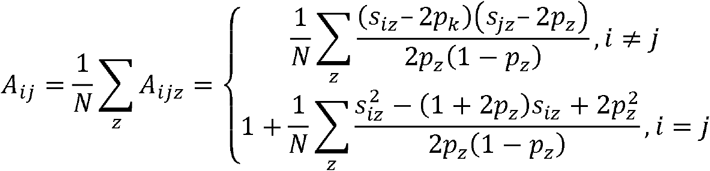

where s_iz_ is the number of copies of the reference allele for SNP z of the individual i, p_z_ is the frequency of the reference allele for the SNP z, and N is the number of SNPs (Yang et al. 2010). The models described above were run using DISSECT (Canela-Xandri et al. 2015).

To deal with multiple measurements per individual for the adult live traits, we used a repeatability model by fitting ID as a random effect to ensure that uncertainty was correctly propagated through all estimations (Mrode 2014). Although DISSECT does not currently have the option to automatically analyse repeated measures, it is possible to modify input files to allow for a repeated measures model (see James et al. (2022) for method details).

For models 1 and 4, we estimated the narrow sense heritability (h^2^) by dividing the additive genetic variance (the variance associated with the GRM or pedigree respectively) by the total estimated phenotypic variance (h^2^_GRM_ and h^2^_ped_ respectively). For models 2 and 3, we estimated three heritabilities: to avoid confusion when comparing with models 1 and 4, we refer to them as h^2^_pop_, the additive genetic variance explained by the full GRM (equivalent to h^2^_GRM_ from model 1); h^2^_kin_, the additive genetic variance explained by the thresholded GRM; and h^2^_pk_, which is estimated as the sum of h^2^_pop_ and h^2^_kin_ (equivalent to h^2^ and h^2^_ped_). For the monogenic traits, we refer to the heritability estimates from models 1 and 3 when only including SNPs on the same chromosome as the causal gene as h^2^_GRM_Chrom_, h^2^_pop_Chrom_, h^2^_kin_Chrom_, and h^2^_pk_Chrom_. Likewise, for the heritability estimates when only including SNPs within 1Mb of the causal gene, we shall refer to them as h^2^_GRM_Region_, h^2^_pop_Region_, h^2^_kin_Chrom_, and h^2^_pk_Chrom_.

We performed log-likelihood ratio tests (LRT) between models 2 and 3 and model 1 at each threshold for each trait. For models where the inclusion of the thresholded GRM was determined to improve the fit, h^2^_pk_ estimates were compared to h^2^_ped_ estimates to determine whether the model including the thresholded GRM was able to estimate h^2^_ped_ accurately, whilst the h^2^_pop_ estimates were compared to h^2^_GRM_ estimates to determine if the h^2^_GRM_ estimates were biased due to the presence of related individuals.

## Results

### Polygenic traits

#### Thresholded GRM models

For the neonate and lamb traits, the inclusion of a thresholded GRM in the model only significantly improved the fit of the model for lamb jaw length for both relationship-thresholded models (model 2, t=0.05 and t=0.1) (Figures 1A, Supplementary Table 2, Supplementary Table 3, Supplementary Table 4).

**Figure 1.**
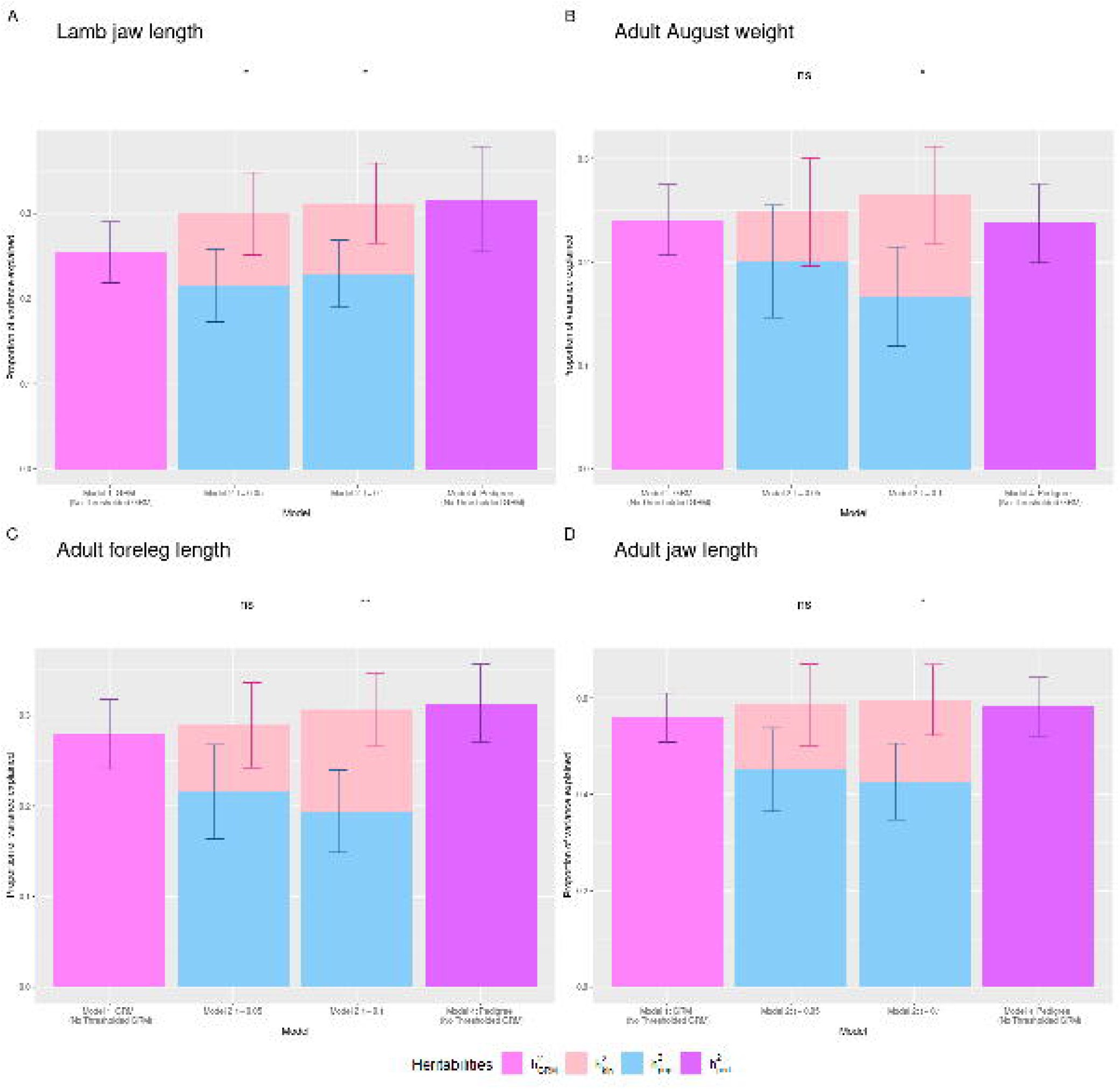
Heritability estimates for **A)** lamb jaw length, **B)** adult August weight, **C)** adult foreleg length and D) adult jaw length for models 1, 2 and 4. For each trait, light purple represents the estimate of h^2^_GRM_ for model 1, blue represents the estimate of h^2^_pop_ for model 2, pink represents the estimate of h^2^_kin_ for model 2, and purple represents the estimate of h^2^_ped_ for model 4. Stars represent the p value when performing loglikelihood ratio tests between models 1 and 2: ns means p > 0.05, * = 0.01 < p < 0.05, ** = p < 0.01.

For lamb jaw length, the h^2^_kin_ estimates were 0.084 and 0.083 for t=0.05 and t=0.1 respectively (S.E. 0.048 and 0.047 respectively) whilst h^2^_pop_ estimates were 0.215 and 0.229 (S.E. 0.043 and 0.039), suggesting that family-associated variance explains 8% of phenotypic variance and 26.5 – 28.1% of the genetic variance underpinning lamb jaw length, depending on the threshold used. For this trait, the inclusion of the MAF-thresholded GRMs (model 3) led to non-convergence of the model. Estimates of h^2^_pk_ (h^2^_pop_ + h^2^_kin_) were higher than that of h^2^_GRM_ for the two relatedness-thresholded models (h^2^_GRM_: 0.254 (S.E. 0.036), h^2^_pk_ t=0.05: 0.300, t=0.1: 0.312), with the h^2^_pk_ estimate for the higher threshold being the highest. The two h^2^_pk_ estimates were similar to the estimate of h^2^_ped_ (0.317 (S.E. 0.062)), with the h^2^_pk_ estimate at t=0.1 falling within the standard error around the estimate of h^2^_ped_. The standard errors around the estimates of h^2^_pop_ overlapped with that of h^2^_GRM_, suggesting that the h^2^_GRM_ estimate for this trait does not suffer from major biases from family-associated affects such as dominance and epistasis (Table 4).

Of the adult traits, the inclusion of a relatedness-thresholded GRM was significant for August weight, foreleg length and jaw length at t=0.1 (Figures 1B, 1C and 1D, Table 4, Supplementary Table 2, Supplementary Table 4). For adult August weight, the estimate of h^2^_kin_ was 0.098 (S.E. 0.047) and the estimate of h^2^_pop_ was 0.167 (S.E. 0.048), suggesting that family-level genetic variance explains 9.8% of the phenotypic variation of this trait and made up 37.1% of the underlying genetic variance. For adult foreleg length, the estimate of h^2^_kin_ was 0.112 (S.E. 0.041) and the estimate of h^2^_pop_ was 0.194 (S.E. 0.045), suggesting that family-level genetic variance explains 11.2% of phenotypic variation and 36.6% of underlying genetic variance for adult foreleg length. For adult jaw length, the estimate of h^2^_kin_ was 0.169 (S.E. 0.074) and the estimate of h^2^_pop_ was 0.427 (S.E. 0.079), suggesting that family-level variance explained 16.9% of the phenotypic variation and 28.4% of the underlying genetic variance for this trait. Estimates of h^2^_pk_ of all three traits fell within the standard errors around their respective h^2^_ped_ estimates, and for adult August weight, the standard error around the estimate of h^2^_pop_overlapped with that of h^2^_GRM_. However, for adult foreleg length and adult jaw length the standard errors around the estimates of h^2^_pop_ and h^2^_GRM_ did not overlap, suggesting that there may be some effect of family-associated variance such as dominance or epistasis biasing the h^2^_GRM_ estimates (Table 4).

The MAF-thresholded models (model 3) did not yield a significant change in the additive genetic variance explained for any trait (Supplementary Table 3, Supplementary Table 4).

#### Monogenic traits

Inclusion of either the relatedness-thresholded GRM (Model 2, t=0.05 and t=0.1) or the MAF-thresholded GRM (Model 3) was not significant for any of the monogenic traits we investigated (Supplementary Table 5, Supplementary Table 6, Supplementary Table 7). These results are somewhat surprising, as dominance is known to play a role in all four of these traits (Dolling 1961; Ryder et al. 1974; Kinsmann 2001; Coltman and Pemberton 2003; Gratten et al. 2007; Gratten et al. 2010; Johnston et al. 2013).

Re-running model 3 focusing on smaller SNP windows (focal chromosome and 1Mb either side of the causal gene) did not improve the model fit for any of the MAF thresholds across any of the traits. For male and female horn type, the regional model was only run for MAF=0.1 and MAF=0.05 due to the fact that less than 10 SNPs remained when applying the more stringent thresholds. For coat colour and coat pattern, the regional model was not run for MAF=0.005 and MAF=0.001 for the same reasons, however the model at the remaining thresholds failed to converge.

Interestingly, for male horn type, coat colour, and coat pattern, estimates of h^2^_GRM_Chrom_ and h^2^_GRM_Region_ from the chromosome and regional models that did run were lower than estimates of h^2^_GRM_ when the whole genome SNP data was used to construct the GRM. The standard error around the estimate of h^2^_GRM_ did not overlap with those of h^2^_GRM_Chrom_ and h^2^_GRM_Region_, suggesting that these estimates are significantly different from each other. For female horn type, h^2^_GRM_Chrom_ and h^2^_GRM_Region_ estimates for the chromosomal and regional models that did run were slightly higher than h^2^_GRM_, though the standard errors around the estimates did overlap.

## Discussion

As far as we are aware, Zaitlen et. al’s method has been used to estimate heritability in populations containing related individuals in humans (Zaitlen et al. 2013), one livestock population (Kokkinias 2022) and two wild populations of birds (Silva et al. 2017). Population structure and sample size both differ between the different studies, as well as the proportion of h^2^_pk_ attributed to h^2^_kin_. Zaitlen et al. (2013) focused on a sample of 38K Icelandic individuals including sibling, half-sibling, parent-offspring, grandparent-grandchild and avuncular relationships, and found that h^2^_kin_ made up 25-67% of h^2^_pk_ estimates across their focal traits. Kokkinias (2022) focused on 5K individuals in a commercial salmon breeding program which contained four breeding lines, with most individuals in a breeding line are closely related (usually either full siblings or half siblings) – they found that h^2^_kin_ made up 0-10% of h^2^_pk_ estimates. Silva et al. focused on both a sample of 700-1400 Norwegian house sparrows and a sample of ∼800 Swedish collared flycatchers, with the sample of house sparrows being more related to each other than the sample of collared flycatchers were. h^2^_kin_ made up 19-100% of h^2^_pk_ in the house sparrow data and 2-74% of h^2^_pk_ estimates in the collared flycatcher data. The Soay sheep population is comprised of related individuals with very complex family structures – both males and females are promiscuous meaning full-siblings are much rarer than half-siblings, and the population is uniformly inbred. For the traits in which inclusion of a thresholded GRM improved model fit, we found that h^2^_kin_ estimates made up 26.5-37.1% of h^2^_pk_ estimates.

It is possible that the difference between these results is because of population relatedness structure; the salmon population was essentially made up of four families and contains no individuals without any relatives, thus the GRM and the thresholded GRM were probably similar. In comparison, the human and bird data comprised a more complex set of relationships resulting in a bigger difference between the GRM and thresholded GRM. Xia et al. (2016) performed a similar analysis to Zaitlen et al. in a population of Scottish humans, but including environmental relationship matrices in the models alongside the thresholded GRM and performed stepwise model selection to determine which combination of matrices resulted in the best model fit for each of their traits. They first did this on a sample of 10K individuals, primarily nuclear families and unrelated individuals, and then on a sample of 20K individuals, made up of the original 10K plus an additional 10K individuals that were mainly related to the original 10K individuals. Of the 16 traits studied, the relatedness-thresholded GRM was retained after model selection for 10 traits when using the 10K data and for 14 when using the 20K data. The authors suggest that the increase in sample size and inclusion of more distant relationships allowed the model to separate the effect of family-associated genetic variance (h^2^_kin_) from family-associated environmental variance for the four traits for which the relatedness-thresholded GRM was retained only when using the 20K data.

We identified three traits that are potentially influenced by family-associated variance. For weight and foreleg length, this effect was only observed in adulthood, whilst for jaw length the effect was found in both lambs and adults.

As jaw length is a skeletal trait, it is measured *post-mortem* and thus there is no overlap between individuals who have a recorded lamb jaw length and those who have a recorded adult jaw length. This suggests that the effect of family-level genetic variance on jaw length is not dependent on any age-related mortality factors such as juvenile survival and is potentially consistent throughout life, affecting both juvenile jaw length and total potential (adult) jaw length.

Weight and foreleg length are live measures, meaning there is an overlap between individuals measured as adults and individuals measured as lambs (and in the case of weight, individuals measured as neonates). In addition, individuals can be caught across multiple years as adults, meaning that adult August weight and adult foreleg length might have repeated measures. There are therefore two main potential reasons as to why our models showed an effect of family-level genetic variance in adulthood but not in juveniles or neonates. Firstly, h^2^ estimates for birth weight, lamb August weight and lamb foreleg length are low (h^2^ < 0.15), so these traits are likely primarily controlled by environmental factors (including maternal effects). Thus, any family-level genetic variation affecting these traits may also be small. Secondly, it is possible that any family-level genetic variation affecting weight and foreleg length only affect the total potential (adult) weight and foreleg length, rather than juvenile weight and foreleg length. This means the effect would only be picked up among individuals who have reached their potential for these traits (i.e. individuals who have stopped growing – adults).

The standard errors around the h^2^_pop_ estimates for adult foreleg length and adult jaw length at t=0.01 did not overlap with those for their respective h^2^_GRM_ estimates, suggesting that any family-associated variance that is influencing these traits is significantly impacting estimates of h^2^_GRM_ when related individuals are included in the analysis. It is interesting that, despite there being a genetic correlation of 0.944 (S.E. 0.019) between adult foreleg length and adult hindleg length (Bérénos et al. 2014), inclusion of a thresholded GRM at t=0.1 is significant for adult foreleg length and not adult hindleg length. However in a recent genome-wide association study of this population, we found two peaks on chromosomes 7 and 9 that were significantly associated with adult foreleg length but not with adult hindleg length (James et al. 2022). It is possible that family-associated variation such as epistasis and dominance are acting upon these two regions, resulting in the inclusion of the thresholded GRM being significant for adult foreleg length but not adult hindleg length. Further investigation, with larger sample sizes that may become available in the future, would shed more light on the commonalities and differences in the determinants of these traits.

Inclusion of either the relatedness-thresholded GRM models (model 2) or the MAF-thresholded models (model 3) did not improve model fit for any of the monogenic traits we investigated. From our results, this model is not well suited for detecting any dominance in monogenic traits. It is possible that this is because the minor allele frequencies of the causal variants are too high to influence the thresholded GRMs – especially the MAF-thresholded GRMs, given that the minor allele frequencies are all higher than our most lenient threshold (Supplementary Table 1). Interestingly, for three of the traits (male horn type, coat colour and coat pattern), estimates of h^2^_GRM_Chrom_ and h^2^_GRM_REgion_ were lower than that of h^2^_GRM_, suggesting that there may be some additive genetic variance influencing these traits located elsewhere in the genome, and that these traits may not be truly monogenic. Further investigation using methods such as regional genomic relationship mapping (Nagamine et al. 2012) or haplotype heritability mapping (Shirali et al. 2018) may therefore provide new insights to the genetic architecture underpinning these traits.

Overall, we have demonstrated that the method proposed by Zaitlen *et al*. (2013) can be useful for estimating heritability in wild population samples containing relatives, as well as showing how the method is affected by different genetic architectures of traits (monogenic versus polygenic). However, we found that the model including the thresholded GRMs was not a significantly better fit for the Soay sheep data when estimating heritability for most of our traits. On the other hand, there are indications have also discovered that some traits previously thought to be monogenic in the Soay sheep may be influenced by genetic variation outwith the causal gene.

## Data Availability Statement

All scripts and data can be found at https://github.com/CaelinnJames/PartioningHeritability_in_SoaySheep

## Acknowledgments

We thank the National Trust for Scotland and Scottish Natural Heritage for permission to work on St Kilda and QinetiQ and Eurest for logistics and other support on the island. We also thank all those who have been involved in the long-term project, including those who helped with field work on the island. We thank the Wellcome Trust Clinical Research Facility Genetics Core in Edinburgh for SNP genotyping.

