## Supplementary Table 1 for "Investigating pedigree- and SNP-associated components of heritability in a wild population of Soay sheep"

| | Causal gene | Chromsome | Gene start bp | Gene end bp | MAF |
|---|---|---|---|---|---|
| Male horn type | RXFP2 | 10 | 29,454,677 | 29,502,617 | 0.471* |
| Female horn type | RXFP2 | 10 | 29,454,677 | 29,502,617 | 0.471* |
| Coat colour | TYRP1 | 2 | 80,602,298 | 80,623,437 | 0.49 |
| Coat pattern | ASIP | 13 | 63,237,431 | 63,242,627 | 0.226** |

Supplementary Table 1: Causal genes, the location of the causal genes (chromsome, start bp and end bp) and minor allele frequencies (MAF) of the monogenic traits.

* Equilibrium frequency

** The self-type coat pattern has been linked to up to five haplotypes which contain any of three recessive variants in the *ASIP* gene. The MAF reported here is the frequency of all five recessive haplotypes combined.
