## Supplementary Table 2 for "Investigating pedigree- and SNP-associated components of heritability in a wild population of Soay sheep"

| | | | Neonate Birth weight | |
|---|---|---|---|---|
| Model 1 | | h2GRM | 0.0504848 | (0.0189423) |
| Model 2 | t=0.05 | h2pop | 0.0310285 | (0.0246469) |
| | | h2kin | 0.0343825 | (0.0282527) |
| | | h2pk | 0.0654110 | |
| | t=0.1 | h2pop | 0.0453195 | (0.0230756) |
| | | h2kin | 0.0100933 | (0.0261491) |
| | | h2pk | 0.0554128 | |
| Model 4 | | h2ped | 0.0782698 | (0.0274258) |

| | | | Lamb | | | | | | | | | |
|---|---|---|---|---|---|---|---|---|---|---|---|---|
| | | | August weight | | Foreleg length | | Hindleg length | | Metacarpal length | | Jaw length | |
| Model 1 | | h2GRM | 0.0836002 | (0.0227376) | 0.1401640 | (0.0250416) | 0.1406110 | (0.0265688) | 0.3198500 | (0.0378536) | 0.2543860 | (0.0357230) |
| Model 2 | t=0.05 | h2pop | 0.0685738 | (0.0310563) | 0.1102290 | (0.0316158) | 0.1165390 | (0.0351703) | 0.2954930 | (0.0471020) | 0.2155440 | (0.0426954) |
| | | h2kin | 0.0263117 | (0.0362078) | 0.0450873 | (0.0326143) | 0.0402105 | (0.0385569) | 0.0420406 | (0.0509113) | 0.0844032 | (0.0484278) |
| | | h2pk | 0.0948855 | | 0.1553163 | | 0.1567495 | | 0.3375336 | | 0.2999472 | |
| | t=0.1 | h2pop | 0.0835192 | (0.0279068) | 0.1182790 | (0.0288547) | 0.1261280 | (0.0318137) | 0.3004040 | (0.0428648) | 0.2292180 | (0.0394293) |
| | | h2kin | 0.0001715 | (0.0318213) | 0.0399467 | (0.0292312) | 0.0297991 | (0.0345896) | 0.0465531 | (0.0482062) | 0.0830081 | (0.0470872) |
| | | h2pk | 0.0836907 | | 0.1582257 | | 0.1559271 | | 0.3469571 | (0.0657066) | 0.3122261 | |
| Model 4 | | h2ped | 0.0920266 | (0.0307689) | 0.1857650 | (0.0351451) | 0.1970670 | (0.0396617) | 0.4068190 | | 0.3167840 | (0.0619316) |

| | | | Adult | | | | | | | | | |
|---|---|---|---|---|---|---|---|---|---|---|---|---|
| | | | August weight | | Foreleg length | | Hindleg length | | Metacarpal length | | Jaw length | |
| Model 1 | | h2GRM | 0.2407890 | (0.0341295) | 0.2789680 | (0.0383702) | 0.4408300 | (0.0408771) | 0.6507510 | (0.0478387) | 0.5597250 | (0.0511207) |
| Model 2 | t=0.05 | h2pop | 0.2007170 | (0.0542305) | 0.2154630 | (0.0520131) | 0.4582100 | (0.0730839) | 0.6552330 | (0.0877115) | 0.4511790 | (0.0866933) |
| | | h2kin | 0.0478639 | (0.0521969) | 0.0735910 | (0.0468383) | 0.0000009 | (0.0700422) | 0.0000010 | (0.0887920) | 0.1347760 | (0.0842002) |
| | | h2pk | 0.2485809 | | 0.2890540 | | 0.4582109 | | 0.6552340 | | 0.5859550 | |
| | t=0.1 | h2pop | **0.1669180** | (0.0478874) | **0.1939620** | (0.0454238) | 0.3804200 | (0.0641476) | 0.6272610 | (0.0753312) | 0.4270150 | (0.0793274) |
| | | h2kin | 0.0982557 | (0.0467192) | 0.1121880 | (0.0407194) | 0.0752228 | (0.0599039) | 0.0303275 | (0.0720732) | 0.1689700 | (0.0735212) |
| | | h2pk | 0.2651737 | | 0.3061500 | | 0.4556428 | | 0.6575885 | | 0.5959850 | |
| Model 4 | | h2ped | 0.2377390 | (0.0382416) | 0.3131680 | (0.0431046) | 0.4447200 | (0.0472902) | 0.6589970 | (0.0582174) | 0.5821470 | (0.0606762) |

(green) Indicates that the LRT gave a p value < 0.05 for the inclusion of the thresholded GRM

**BOLD** Indicates that the h2pop estimate does not fall within the standard error of the h2GRM estimate

Supplementary Table 2: h2GRM, h2pop, h2kin, h2pk and h2ped estimates for each polygenic trait for models 1, 2 and 4 with their respective standard errors. h2pk is calculated as the sum of h2kin and h2pop. Green highlight indicates that the LRT gave a p value < 0.05 for the inclusion of the thresholded GRM, whilst bolded h2pk estimates indicate that the h2pop estimate does not fall within the standard error of the respective h2GRM estimate.
