## Supplementary Table 3 for "Investigating pedigree- and SNP-associated components of heritability in a wild population of Soay sheep"

| | | | Neonate | | Lamb | | | | | | | | | |
|---|---|---|---|---|---|---|---|---|---|---|---|---|---|---|
| | | | Birth weight | | August weight | | Foreleg length | | Hindleg length | | Metacarpal length | | Jaw length | |
| Model 1 | | h2GRM | 0.0504848 | (0.0189423) | 0.0836002 | (0.0227376) | 0.1401640 | (0.0250416) | 0.1406110 | (0.0265688) | 0.3198500 | (0.0378536) | 0.2543860 | (0.0357230) |
| Model 3 | t=0.1 | h2pop | 0.0887579 | (0.0355111) | 0.1131290 | (0.0421449) | 0.1838270 | (0.0447654) | 0.2002460 | (0.0491091) | 0.3570040 | (0.0644263) | 0.2922570 | (0.0582801) |
| | | h2kin | 0.0000009 | (0.0319769) | 0.0000009 | (0.0385133) | 0.0000009 | (0.0388257) | 0.0000009 | (0.0445024) | 0.0000009 | (0.0555167) | 0.0000009 | (0.0502495) |
| | | h2pk | 0.0887588 | | 0.1131299 | | 0.1838279 | | 0.2002469 | | 0.3570049 | | 0.2922579 | |
| | t=0.05 | h2pop | 0.0649126 | (0.0256225) | 0.0919911 | (0.0307153) | 0.1600630 | (0.0333593) | 0.1588910 | (0.0354883) | 0.3155620 | (0.0486348) | 0.2714230 | (0.0453895) |
| | | h2kin | 0.0000009 | (0.0197049) | 0.0000009 | (0.0246078) | 0.0000009 | (0.0241801) | 0.0000009 | (0.0271960) | 0.0035804 | (0.0350481) | 0.0000009 | (0.0338638) |
| | | h2pk | 0.0649135 | | 0.0919920 | | 0.1600639 | | 0.1588919 | | 0.3191424 | | 0.2714239 | |
| | t=0.01 | h2pop | 0.0476112 | (0.0194963) | 0.0759578 | (0.0232446) | 0.1375290 | (0.0257127) | 0.1416860 | (0.0276798) | 0.3207590 | (0.0397298) | 0.2562250 | (0.0370333) |
| | | h2kin | 0.0039096 | (0.0091319) | 0.0179776 | (0.0141600) | 0.0043647 | (0.0105374) | 0.0000010 | (0.0120848) | 0.0000010 | (0.0166571) | 0.0000010 | (0.0157298) |
| | | h2pk | 0.0515208 | | 0.0939354 | | 0.1418937 | | 0.1416870 | | 0.3207600 | | 0.2562260 | |
| | t=0.005 | h2pop | 0.0447857 | (0.0188276) | 0.0794582 | (0.0229079) | 0.1371820 | (0.0252384) | 0.1411110 | (0.0271742) | 0.3197130 | (0.0387698) | 0.2546130 | (0.0363830) |
| | | h2kin | 0.0091967 | (0.0082832) | 0.0092595 | (0.0105304) | 0.0060356 | (0.0088084) | 0.0000010 | (0.0091289) | 0.0000010 | (0.0120349) | 0.0000010 | (0.0113618) |
| | | h2pk | 0.0539824 | | 0.0887177 | | 0.1432176 | | 0.1411120 | | 0.3197140 | | 0.2546140 | |
| | t=0.001 | h2pop | 0.0505879 | (0.0189987) | 0.0858074 | (0.0231565) | 0.1403890 | (0.0251294) | 0.1433450 | (0.0269713) | 0.3191210 | (0.0380116) | 0.2531430 | (0.0357344) |
| | | h2kin | 0.0000010 | (0.0035382) | 0.0000009 | (0.0060963) | 0.0000010 | (0.0038095) | 0.0000010 | (0.0063960) | 0.0000010 | (0.0067004) | 0.0000010 | (0.0060550) |
| | | h2pk | 0.0505889 | | 0.0858083 | | 0.1403900 | | 0.1433460 | | 0.3191220 | | 0.2531440 | |
| Model 4 | | h2ped | 0.0782698 | (0.0274258) | 0.0920266 | (0.0307689) | 0.1857650 | (0.0351451) | 0.1970670 | (0.0396617) | 0.4068190 | (0.0657066) | 0.3167840 | (0.0619316) |

| | | | Adult | | | | | | | | | |
|---|---|---|---|---|---|---|---|---|---|---|---|---|
| | | | August weight | | Foreleg length | | Hindleg length | | Metacarpal length | | Jaw length | |
| Model 1 | | h2GRM | 0.2407890 | (0.0341295) | 0.2789680 | (0.0383702) | 0.4408300 | (0.0408771) | 0.6507510 | (0.0478387) | 0.5597250 | (0.0511207) |
| Model 3 | t=0.1 | h2pop | 0.2653940 | (0.0684382) | 0.2993080 | (0.0652254) | 0.4451120 | (0.0903141) | 0.6851390 | (0.1121900) | 0.5783910 | (0.1056780) |
| | | h2kin | 0.0000009 | (0.0612993) | 0.0000009 | (0.0537638) | 0.0000010 | (0.0823708) | 0.0000009 | (0.1071730) | 0.0000009 | (0.0948522) |
| | | h2pk | 0.2653949 | | 0.2993089 | | 0.4451130 | | 0.6851399 | | 0.5783919 | |
| | t=0.05 | h2pop | 0.2430760 | (0.0497712) | 0.2689810 | (0.0487087) | 0.4185690 | (0.0659350) | 0.6770240 | (0.0813763) | 0.5824480 | (0.0785137) |
| | | h2kin | 0.0000009 | (0.0384626) | 0.0108812 | (0.0331931) | 0.0231210 | (0.0545600) | 0.0000009 | (0.0718124) | 0.0000009 | (0.0625187) |
| | | h2pk | 0.2430769 | | 0.2798622 | | 0.4416900 | | 0.6770249 | | 0.5824489 | |
| | t=0.01 | h2pop | 0.2264140 | (0.0374344) | 0.2752630 | (0.0404486) | 0.4465800 | (0.0469805) | 0.6587400 | (0.0542465) | 0.5741510 | (0.0560763) |
| | | h2kin | 0.0138312 | (0.0182794) | 0.0034573 | (0.0135985) | 0.0000010 | (0.0226367) | 0.0000009 | (0.0296998) | 0.0000009 | (0.0240457) |
| | | h2pk | 0.2402452 | | 0.2787203 | | 0.4465810 | | 0.6587409 | | 0.5741519 | |
| | t=0.005 | h2pop | 0.2265440 | (0.0357928) | 0.2810670 | (0.0396063) | 0.4461100 | (0.0441867) | 0.6523690 | (0.0507328) | 0.5696410 | (0.0533266) |
| | | h2kin | 0.0127770 | (0.0139642) | 0.0000009 | (0.0091444) | 0.0000010 | (0.0161608) | 0.0000010 | (0.0188489) | 0.0000009 | (0.0155367) |
| | | h2pk | 0.2393210 | | 0.2810679 | | 0.4461110 | | 0.6523700 | | 0.5696419 | |
| | t=0.001 | h2pop | 0.2485630 | (0.0353771) | 0.2780550 | (0.0385074) | 0.4420260 | (0.0420207) | 0.6444640 | (0.0484995) | 0.5636970 | (0.0514328) |
| | | h2kin | 0.0000009 | (0.0080807) | 0.0010970 | (0.0049754) | 0.0000010 | (0.0085612) | 0.0000010 | (0.0051735) | 0.0000009 | (0.0036250) |
| | | h2pk | 0.2485639 | | 0.2791520 | | 0.4420270 | | 0.6444650 | | 0.5636979 | |
| Model 4 | | h2ped | 0.2377390 | (0.0382416) | 0.3131680 | (0.0431046) | 0.4447200 | (0.0472902) | 0.6589970 | (0.0582174) | 0.5821470 | (0.0606762) |

Supplementary Table 3: h2GRM, h2pop, h2kin, h2pk and h2ped estimates for each polygenic trait for models 1, 3 and 4 with their respective standard errors. h2pk is calculated as the sum of h2kin and h2pop.
