## Supplementary Table 4 for "Investigating pedigree- and SNP-associated components of heritability in a wild population of Soay sheep"

| | | Neonate | Lamb | | | | |
|---|---|---|---|---|---|---|---|
| | | Birth weight | August weight | Foreleg length | Hindleg length | Metacarpal length | Jaw length |
| | t=0.05 | 0.09867 | 0.2286 | 0.08066 | 0.1525 | 0.2017 | 0.04067 |
| Model 2 | t=0.1 | 0.3461 | 0.4978 | 0.0813 | 0.1976 | 0.159 | 0.03831 |
| | MAF=0.1 | NA | NA | NA | NA | NA | NA |
| | MAF=0.05 | NA | NA | NA | NA | 0.4605 | NA |
| | MAF=0.01 | 0.3194 | 0.1031 | 0.3295 | NA | NA | NA |
| | MAF=0.005 | 0.1012 | 0.1945 | 0.2323 | NA | NA | NA |
| Model 3 | MAF=0.001 | NA | NA | NA | NA | NA | NA |

| | | Adult | | | | |
|---|---|---|---|---|---|---|
| | | August weight | Foreleg length | Hindleg length | Metacarpal length | Jaw length |
| | t=0.05 | 0.1643 | 0.06552 | NA | NA | 0.05255 |
| Model 2 | t=0.1 | 0.01435 | 0.003718 | 0.1067 | 0.3315 | 0.01074 |
| | MAF=0.1 | NA | NA | NA | NA | NA |
| | MAF=0.05 | NA | 0.3687 | 0.3361 | NA | NA |
| | MAF=0.01 | 0.211 | 0.3964 | NA | NA | NA |
| | MAF=0.005 | 0.1611 | NA | NA | NA | NA |
| Model 3 | MAF=0.001 | NA | 0.4046 | NA | NA | NA |

Supplementary Table 4: p values for the LRT for the inclusion of the thresholded GRMs for each threshold for models 2 and 3 for each polygenic trait. "NA" means that the model did not converge for that trait and threshold - we consider this to be equivelent to a p value of 1. Green highlights cells with a p value < 0.05
