## Supplementary Table 5 for "Investigating pedigree- and SNP-associated components of heritability in a wild population of Soay sheep"

| | | | Male horn type | | Female horn type | | Coat colour | | Coat pattern | |
|---|---|---|---|---|---|---|---|---|---|---|
| Model 1 | | h2GRM | 0.3518910 | (0.0269923) | 0.2282130 | (0.0267518) | 0.6132300 | (0.0127196) | 0.3499710 | (0.0161968) |
| | | h2pop | 0.3177530 | (0.0345399) | 0.2174220 | (0.0369595) | 0.6054230 | (0.0151641) | 0.3587990 | (0.0194481) |
| | | h2kin | 0.0583169 | (0.0350713) | 0.0151738 | (0.0366140) | 0.0130067 | (0.0125519) | 0.0000010 | (0.0160213) |
| | t=0.05 | h2pk | 0.3760699 | | 0.2325958 | | 0.6184297 | | 0.3588000 | |
| | | h2pop | 0.3371430 | (0.0319960) | 0.2247330 | (0.0338976) | 0.6064820 | (0.0146091) | 0.3532080 | (0.0182692) |
| | | h2kin | 0.0288222 | (0.0315916) | 0.0052941 | (0.0323712) | 0.0116538 | (0.0113208) | 0.0000010 | (0.0140086) |
| Model 2 | t=0.1 | h2pk | 0.3659652 | | 0.2300271 | | 0.6181358 | | 0.3532090 | |
| Model 4 | | h2ped | 0.3905800 | (0.0369544) | 0.2387470 | (0.0313989) | 0.6529220 | (0.0189878) | 0.3333320 | (0.0222770) |

Supplementary Table 5: h2GRM, h2pop, h2kin, h2pk and h2ped estimates for each monogenic trait for models 1, 2 and 4 with their respective standard errors. h2pk is calculated as the sum of h2kin and h2pop.
