## Supplementary Table 6 for "Investigating pedigree- and SNP-associated components of heritability in a wild population of Soay sheep"

| | | | Male horn type | | | | | | Female horn type | | | | | |
|---|---|---|---|---|---|---|---|---|---|---|---|---|---|---|
| | | | Whole genome | | Chromosome | | 1Mb either side of gene | | Whole genome | | Chromosome | | 1Mb either side of gene | |
| Model 1 | | h2GRM | 0.3518910 | (0.0269923) | 0.2505240 | (0.0255399) | 0.1294440 | (0.0236076) | 0.2282130 | (0.0267518) | 0.2555530 | (0.0273251) | 0.1993230 | (0.0304379) |
| Model 3 | t=0.1 | h2pop | 0.2919600 | (0.0441054) | 0.3011640 | (0.0449449) | 0.1231030 | (0.0266039) | 0.2498050 | (0.0491979) | 0.3736430 | (0.0560876) | 0.2051340 | (0.0326240) |
| | | h2kin | 0.0648018 | (0.0401937) | 0.0000009 | (0.0373498) | 0.0219039 | (0.0225205) | 0.0000010 | (0.0422002) | 0.0000008 | (0.0498471) | 0.0000010 | (0.0421791) |
| | | h2pk | 0.3567618 | | 0.3011649 | | 0.1450069 | | 0.2498060 | | 0.3736438 | | 0.2051350 | |
| | t=0.05 | h2pop | 0.3526700 | (0.0351791) | 0.2915060 | (0.0338265) | 0.1305370 | (0.0241229) | 0.2308140 | (0.0371107) | 0.3068430 | (0.0381716) | 0.2025420 | (0.0312843) |
| | | h2kin | 0.0000010 | (0.0256721) | 0.0000010 | (0.0227651) | 0.0000011 | (0.0103728) | 0.0000010 | (0.0263063) | 0.0000009 | (0.0256807) | 0.0000010 | (0.0231875) |
| | | h2pk | 0.3526710 | | 0.2915070 | | 0.1305381 | | 0.2308150 | | 0.3068439 | | 0.2025430 | |
| | t=0.01 | h2pop | 0.3501960 | (0.0283281) | 0.2538920 | (0.0264127) | | | 0.2308020 | (0.0284570) | 0.2607610 | (0.0284598) | | |
| | | h2kin | 0.0015435 | (0.0111964) | 0.0000010 | (0.0054208) | | | 0.0000010 | (0.0107354) | 0.0000010 | (0.0057006) | | |
| | | h2pk | 0.3517395 | | 0.2538930 | | | | 0.2308030 | | 0.2607620 | | | |
| | t=0.005 | h2pop | 0.3523240 | (0.0276263) | 0.2518410 | (0.0259569) | | | 0.2261640 | (0.0275762) | 0.2628470 | (0.0282689) | | |
| | | h2kin | 0.0000010 | (0.0076378) | 0.0000010 | (0.0038393) | | | 0.0025333 | (0.0077549) | 0.0000010 | (0.0051967) | | |
| | | h2pk | 0.3523250 | | 0.2518420 | | | | 0.2286973 | | 0.2628480 | | | |
| | t=0.001 | h2pop | 0.3520460 | (0.0271412) | 0.2535350 | (0.0258705) | | | 0.2281370 | (0.0269024) | 0.2613130 | (0.0279011) | | |
| | | h2kin | 0.0000010 | (0.0032549) | 0.0000010 | (0.0033962) | | | 0.0004658 | (0.0027910) | 0.0000010 | (0.0049876) | | |
| | | h2pk | 0.3520470 | | 0.2535360 | | | | 0.2286028 | | 0.2613140 | | | |
| Model 4 | | h2ped | 0.3905800 | (0.0369544) | 0.3905800 | (0.0369544) | 0.3905800 | (0.0369544) | 0.2387470 | (0.0313989) | 0.2387470 | (0.0313989) | 0.2387470 | (0.0313989) |

| | | | Coat colour | | | | | | Coat pattern | | | | | |
|---|---|---|---|---|---|---|---|---|---|---|---|---|---|---|
| | | | Whole genome | | Chromosome | | 1Mb either side of gene | | Whole genome | | Chromosome | | 1Mb either side of gene | |
| Model 1 | | h2GRM | 0.6132300 | (0.0127196) | 0.5229850 | (0.0160838) | Failed to converge | | 0.3499710 | (0.0161968) | 0.2433330 | (0.0192692) | Failed to converge | |
| Model 3 | t=0.1 | h2pop | 0.6666610 | (0.0274822) | 0.5728020 | (0.0321004) | | | 0.4370840 | (0.0285590) | 0.2884420 | (0.0286982) | | |
| | | h2kin | 0.0000009 | (0.0281649) | 0.0000011 | (0.0350071) | Failed to converge | | 0.0000008 | (0.0277818) | 0.0000011 | (0.0213478) | Failed to converge | |
| | | h2pk | 0.6666619 | | 0.5728031 | | | | 0.4370848 | | 0.2884431 | | | |
| | t=0.05 | h2pop | 0.6406340 | (0.0199443) | 0.5640030 | (0.0255429) | | | 0.3860980 | (0.0214723) | 0.2544570 | (0.0232076) | | |
| | | h2kin | 0.0000010 | (0.0177994) | 0.0000011 | (0.0246360) | Failed to converge | | 0.0000009 | (0.0168216) | 0.0000011 | (0.0098138) | Failed to converge | |
| | | h2pk | 0.6406350 | | 0.5640041 | | | | 0.3860989 | | 0.2544581 | | | |
| | t=0.01 | h2pop | 0.6232530 | (0.0142494) | 0.5370430 | (0.0181533) | | | 0.3544790 | (0.0170833) | 0.2595920 | (0.0210876) | | |
| | | h2kin | 0.0000011 | (0.0071022) | 0.0000012 | (0.0114757) | Failed to converge | | 0.0000010 | (0.0065411) | 0.0000011 | (0.0062941) | Failed to converge | |
| | | h2pk | 0.6232541 | | 0.5370442 | | | | 0.3544800 | | 0.2595931 | | | |
| | t=0.005 | h2pop | 0.6207620 | (0.0136306) | 0.5291660 | (0.0170116) | | | 0.3508820 | (0.0166385) | 0.2514000 | (0.0201258) | | |
| | | h2kin | 0.0000011 | (0.0052087) | 0.0000012 | (0.0073172) | | | 0.0000010 | (0.0043203) | 0.0000011 | (0.0048451) | | |
| | | h2pk | 0.6207631 | | 0.5291672 | | | | 0.3508830 | | 0.2514011 | | | |
| | t=0.001 | h2pop | 0.6177950 | (0.0129931) | 0.5235020 | (0.0161557) | | | 0.3488810 | (0.0162390) | 0.2437710 | (0.0193240) | | |
| | | h2kin | 0.0000011 | (0.0016679) | 0.0000012 | (0.0018069) | | | 0.0000010 | (0.0015172) | 0.0000011 | (0.0026712) | | |
| | | h2pk | 0.6177961 | | 0.5235032 | | | | 0.3488820 | | 0.2437721 | | | |
| Model 4 | | h2ped | 0.6529220 | (0.0189878) | 0.6529220 | (0.0189878) | 0.6529220 | (0.0189878) | 0.3333320 | (0.0222770) | 0.3333320 | (0.0222770) | 0.3333320 | (0.0222770) |

Supplementary Table 6: h2GRM, h2pop, h2kin, h2pk and h2ped estimates for each polygenic trait for models 1, 3 and 4 with their respective standard errors. h2pk is calculated as the sum of h2kin and h2pop.

Grey indicates that the model was not ran due to an insufficient number of SNPs remaining after filtering to calculate the GRM from
