## Supplementary Table 7 for "Investigating pedigree- and SNP-associated components of heritability in a wild population of Soay sheep"

| | | Male horn type | | | Female horn type | | |
|---|---|---|---|---|---|---|---|
| | | Whole genome | Chromosome | 1Mb either side of gene | Whole genome | Chromosome | 1Mb either side of gene |
| | t=0.05 | 0.05253 | Not run | Not run | 0.3396 | Not run | Not run |
| Model 2 | t=0.1 | 0.1866 | Not run | Not run | 0.4342 | Not run | Not run |
| | MAF=0.1 | 0.05662 | NA | 0.07261 | NA | NA | NA |
| | MAF=0.05 | NA | NA | NA | NA | NA | NA |
| | MAF=0.01 | 0.4473 | NA | Not run | NA | NA | Not run |
| | MAF=0.005 | NA | NA | Not run | 0.3603 | NA | Not run |
| Model 3 | MAF=0.001 | NA | NA | Not run | 0.4267 | NA | Not run |

| | | Coat colour | | | Coat pattern | | |
|---|---|---|---|---|---|---|---|
| | | Whole genome | Chromosome | 1Mb either side of gene | Whole genome | Chromosome | 1Mb either side of gene |
| | t=0.05 | 0.1502 | Not run | Not run | NA | NA | Not run |
| Model 2 | t=0.1 | 0.1486 | Not run | Not run | NA | NA | Not run |
| | MAF=0.1 | NA | NA | NA | NA | NA | NA |
| | MAF=0.05 | NA | NA | NA | NA | NA | NA |
| | MAF=0.01 | NA | NA | NA | NA | NA | NA |
| | MAF=0.005 | NA | NA | Not run | NA | NA | Not run |
| Model 3 | MAF=0.001 | NA | NA | Not run | NA | NA | Not run |

Supplementary Table 7: p values for the LRT for the inclusion of the thresholded GRMs for each threshold for models 2 and 3 for each monogenic trait. "NA" means that the model did not converge for that trait and threshold - we consider this to be equivelent to a p value of 1. "Not run" means we did not run that model.
